# DeepMAsED: Evaluating the quality of metagenomic assemblies

**DOI:** 10.1101/763813

**Authors:** Mateo Rojas-Carulla, Ruth E. Ley, Bernhard Schölkopf, Nicholas D. Youngblut

## Abstract

**Motivation/background:** Methodological advances in metagenome assembly are rapidly increasing in the number of published metagenome assemblies. However, identifying misassemblies is challenging due to a lack of closely related reference genomes that can act as pseudo ground truth. Existing reference-free methods are no longer maintained, can make strong assumptions that may not hold across a diversity of research projects, and have not been validated on large scale metagenome assemblies.

**Results:** We present DeepMAsED, a deep learning approach for identifying misassembled contigs without the need for reference genomes. Moreover, we provide an *in silico* pipeline for generating large-scale, realistic metagenome assemblies for comprehensive model training and testing. DeepMAsED accuracy substantially exceeds the state-of-the-art when applied to large and complex metagenome assemblies. Our model estimates close to a 5% contig misassembly rate in two recent large-scale metagenome assembly publications.

**Conclusions:** DeepMAsED accurately identifies misassemblies in metagenome-assembled contigs from a broad diversity of bacteria and archaea without the need for reference genomes or strong modelling assumptions. Running DeepMAsED is straight-forward, as well as is model re-training with our dataset generation pipeline. Therefore, DeepMAsED is a flexible misassembly classifier that can be applied to a wide range of metagenome assembly projects.

**Availability:** DeepMAsED is available from GitHub at https://github.com/leylabmpi/DeepMAsED.

## 1 Introduction

The rapid development of affordable DNA sequencing technologies has led to an explosion of the number of prokaryotic genomes assembled from metagenomes (Pasolli *et al.*, 2019; Almeida *et al.*, 2019). Genomes are typically assembled from short DNA sequences using assemblers such as MEGAHIT (Li *et al.*, 2015) and MetaSPAdes (Nurk *et al.*, 2017). These assemblers receive as input a collection of sequenced DNA reads originating from a large number of organisms, and aim to group them into overlapping sections of DNA. This process results in a *contig*, a consensus region of DNA of variable length. A Metagenome-Assembled Genome (MAG) is a collection of such contigs which are estimated to have originated from the same strain. However, evaluating the quality of assembled contigs remains a difficult problem. In contig regions with large *coverage*, that is, where a large number of reads overlap the given region in the contig, large consensus is an indicator of quality, but how to aggregate such findings into a quality score for a full contig remains an open question.

Most methods that assess the quality of assemblies require the availability of reference genomes: a set of curated genomes that the assemblies can be compared against. Such methods include metaQUAST (Mikheenko *et al.*, 2015), which uses mapping of contigs to one or more references to infer *extensive misassemblies* such as inversions, relocations, and interspecies translocations. Nonetheless, several newly discovered MAGs represent novel organisms that are distantly related to organisms represented in the genomic databases, and this makes selecting a valid reference often impossible. For instance, sometimes the closest known relative belongs to a different bacterial phylum, which implies very deep evolutionary divergence (Pasolli *et al.*, 2019). Conversely, because species of Bacteria can include strains with large genomic differences, true genomic variation among strains can be misinterpreted as misassemblies.

The assembly of full or partial genomes from metagenomes would thus benefit from methods *without* reference requirements, but few such methods are available. The most widely-used method is CheckM (Parks *et al.*, 2015), which provides two sub-scores commonly used for quality evaluations: *completeness* and *contamination.* These measures are based on counts of specific loci, and thus do not evaluate genome assembly at the level of individual contigs. Still, large scale MAG datasets such as Almeida *et al.* (2019) and Pasolli *et al.* (2019) use thresholds for completeness and contamination to screen for assembly quality.

Two predominant reference-free methods for evaluating contig misassembly are ALE (Clark *et al.*, 2013) and SuRankCo (Kuhring *et al.*, 2015). ALE makes probabilistic assumptions about the genomes and provides a likelihood score for each nucleotide in an assembled contig, but does not provide a quality estimate at the contig level. Moreover, it is tailored for genomic and not metagenomic assemblies. SuRankCo makes fewer assumptions about the genomes and the assemblers by utilizing a random forest classifier on a set of hand-crafted features to predict a quality estimate for the input contig. Crucially, neither ALE nor SuRankCo are currently maintained, and compatibility issues also hamper their use.

### 1.1 Contributions

Here, we introduce DeepMAsED^1^ (Deep Metagenome Assembly Error Detection), a machine learning system for quality evaluation of metagenomic assemblies at the contig level. The goal of DeepMAsED is to predict the *extensive misassembly* label from metaQUAST *without* the availability of reference genomes. The contributions of our paper can be summarized as follows.

i. We propose a data-generation pipeline for training and for rigorous evaluation. Given an initial set of known genomes, we generate a dataset 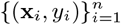, where x_*i*_ represents a raw contig sequence augmented with additional features, and *y_i_* is a binary target indicating whether metaQUAST labels the corresponding contig as misassembled.
ii. We introduce DeepMAsED, a deep convolutional network for assessing assembly quality at the contig level trained on raw sequence and read-count features which makes no explicit assumptions about the genomes or the assemblers. We compare against and significantly outperform ALE on the generated datasets. Moreover, the weights from the fully trained system require a modest 1.3 MB of storage. The challenging evaluation procedure should ensure that DeepMAsED generalizes to a large variety of research projects.
iii. We evaluate DeepMAsED on a subset of two large-scale assembly benchmarks from Almeida *et al.* (2019) and Pasolli *et al.* (2019). An estimate from DeepMASeD finds 5% misassembled contigs in Pasolli *et al.* (2019) and Almeida *et al.* (2019). Moreover, we show that CheckM scores, which were used for assessing assembly quality, are poor indicators of DeepMAsED predictions.

## 2 Materials and Methods

In metagenomic analysis, one principal goal of a Next Generation Sequencing (NGS) assembler is to recover genomes from reads obtained from DNA sequencing. The assembly process results in a set of *contigs*, which are DNA sequences of variable length. To build these outputs, assemblers seek overlapping reads at given positions and output a nucleotide based on the consensus from these reads. The assembly problem is more difficult for metagenomes relative to single organism genome-assemblies due to the higher levels of sequence complexity and high prevalence of highly similar genomic regions (e.g., rRNA operons) among closely related organisms, which can often result in incorrectly assembled chimeric contigs. Our goal is to build a machine learning system that receives a contig and the read count information at each position, and outputs a quality estimate. A higher estimate from DeepMAsED indicates a higher likelihood that the corresponding contig is misassembled according to metaQUAST, a reference based method. First, we describe in more detail the features and target of our model in Section 2.1. We then describe the data generation pipeline in Section 2.2 and introduce the architecture of DeepMAsED in Section 2.3.

### 2.1 Input features and target

A contig is a sequence of nucleotides s ∈ {A, C, T, G}^*L*^, where the length *L* is variable. At each position *j* ∈ {1,…, *L*} in the sequence, we include features from two categories: *raw sequence* features and *read count* features.

*Raw sequence* features are a one-hot encoding of the nucleotide at position *j*: a four dimensional binary vector o^*j*^ equal to one at the position of the nucleotide.

*Read count* features are obtained from mapping the reads back to the assembled contigs using Bowtie2 (Langmead and Salzberg, 2012). Let *C^j^* be the coverage at position *j*, that is, the number of reads that map to position j during the assembly process. Among these reads, let 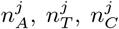 and 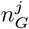 be the count of each nucleotide respectively, with 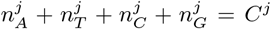. Moreover, let *d^j^* be the number of discordant reads and *v^j^* the number of SNPs at position *j*. Then the read count features correspond to the vector

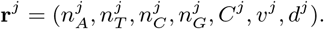

The vector x^*j*^ = (**o**^*j*^, **r**^*j*^) of size *p* =11 obtained by concatenating the raw sequence and read count features is the input feature vector for DeepMAsED at position *j* ∈ {1,…, *L*}, which indicates sources of information about the contig: the raw sequence data, the coverage of the position and the agreement between the corresponding reads.

As an example, assume that the nucleotide at position *j* in the an assembled contig is A. Moreover, assume that ten reads mapped to this position, with six having an A and four having a T at this position, and one read was discordant. The number of SNPs then equals two. Then the feature vector would be

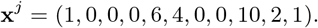

#### Fixed input size and normalization

The convolutional network requires a fixed input contig length, which we set to *L* = 10 000. Shorter contigs are padded with zeroes, while longer contigs are split in non-overlapping chunks of size *L*. An input contig is therefore represented by a matrix x of shape *L × p*, where *L* = 10 000 and *p* = 11. To compute an aggregate score for contigs longer that *L*, we compute a score 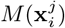 for each non-overlapping sub-sequence *j* of size *L*, and return the average of these scores.

We normalize each non-binary dimension in x as follows. The training dataset 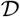 is a matrix **X** of dimension *n × L × p*, where *n* is the number of contigs available in the training set. We reshape **X** into a (*nL*) × *p* matrix by pooling the position dimension, and compute the mean and standard deviation for each of the *p* — 4 read-count features. We then standardize the corresponding dimensions to have mean zero and variance one. The means and standard deviations computed with the training set are stored and used for testing at a later stage.

### 2.2 Generating the synthetic datasets

Training and evaluating DeepMAsED requires a dataset of contigs, read counts and metaQUAST quality labels 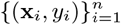. This Section describes the data generation pipeline which consists of i) selecting genomes and simulating genome abundances in several metagenomes, ii) generating reads from these metagenomes in accordance with genome abundances, iii) assembling these reads using standard metagenome assemblers, iv) comparing the output of the assembly (contigs) to the ground truth genomes in order to get ground truth misassembly classifications and v) mapping the reads to each contig in order to generate features at each contig position. The data generation pipeline is summarized in Figure 1.

**Figure 1:**
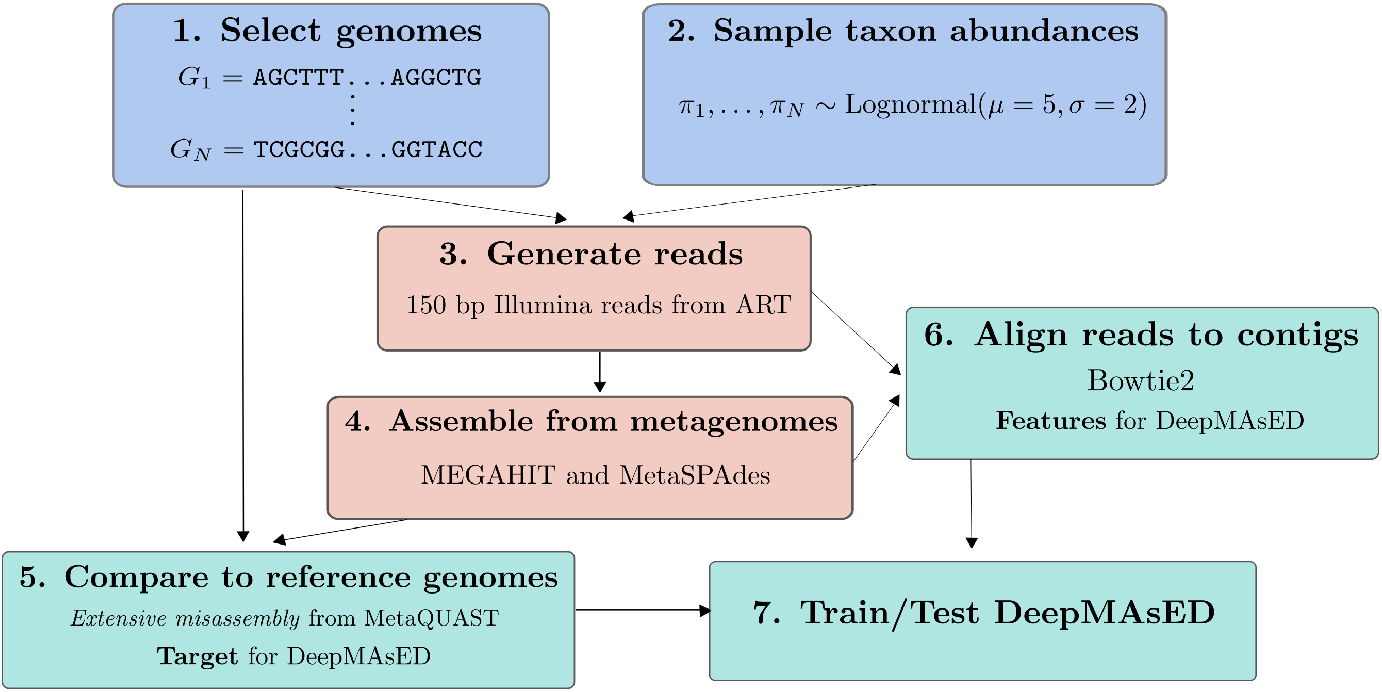
Summary of our data generation pipeline. Given an initial set of *N* genomes, this process results in a dataset 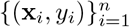, where x_*i*_ is an assembled contig with features described in Section 2.1, and *y_i_* is the corresponding misassembly label from MetaQUAST.

#### Create metagenomes

We generate a dataset 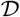 of *N* = 1000 genomes for training and 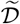 of *Ñ* = 100 non-overlapping genomes for testing. The genomes are randomly selected species representatives from the Genome Taxonomy Database (GTDB) (Parks *et al.*, 2018) after filtering out genomes with a CheckM-estimated completeness of < 50% and contamination ≥ 5%. The genomes consist of both Bacteria and Archaea, with 40 phyla represented. The average estimated completeness and contamination are 98.8 ± 1.85% and 0.73 ± 0.86% respectively, where standard deviation is indicated. These estimates do not substantially differ between the train and test partitions, see Figure S3. The corresponding NCBI accessions are provided in the Supplement.

We utilized MGSIM (Youngblut, 2019) to create several metagenomic communities in which the abundance of each genome is randomly sampled in order to prevent over-fitting to a particular abundance distribution. The training and test sets contain 30 and 25 replicate metagenomes, respectively. We sample taxon abundances for the *N* training genomes from a lognormal distribution *π*_1_,…, *π_N_* ~ Lognormal(*μ, σ*), where *μ* = 5 and *σ* = 2, and randomly permute the obtained abundances in each of the 30 metagenomes. The same procedure is carried out with the *Ñ* genomes in the test set.

#### Generate reads

MGSIM utilizes ART (v2016.06.05) (Huang *et al.*, 2011) for read simulation. We simulated 10^6^ Illumina HiSeq2500 150 base-pair (bp) paired-end reads per metagenome, with the ART-defined default error distribution. This sequencing depth produces a realistic sample coverage of 0.3 to 0.4 per sample, as estimated by Nonpareil (Rodriguez *et al.*, 2018).

#### Assemble genomes from metagenomes

For each obtained metagenome, we provide the simulated reads as input to one of two widely used assemblers: MEGAHIT (v1.2.7) (Li *et al.*, 2015) and MetaSPAdes (v3.13.1) (Nurk *et al.*, 2017). The output of the assembly process are *contigs* of variable length. We filtered out all contigs shorter than 1000 bp. For reference, Figure S1 shows the distribution of log lengths of the assembled contigs for both technologies on the training and test sets. The total number of assembled contigs in the training set is *n* = 418 641 in the training set and *ñ* = 559 237 in the test set.

#### Identify true misassemblies

The ground truth genomes used for metagenome simulation are given for reference-based identification of misassemblies via metaQUAST (v5.0.2) (Mikheenko *et al.*, 2015). The *extensive misassembly* label from metaQUAST is used as a target for DeepMAsED. This umbrella term considers intra-genome *inversions, translocations*, *relocations*, or *interspecies translocations* in the generated data. The distribution of these errors differs slightly between the training and test datasets, see Figure 2, which is likely due to a higher fraction of closely related genomes in the training dataset and thus more chimeric contigs composed of similar genomic regions from multiple organisms. The average nucleotide identity (ANI) distribution for the genomes in the training and test datasets is given in Figure S2.

#### Compute features

We use Bowtie2 (v2.3.5) (Langmead and Salzberg, 2012) to map the simulated reads back to the metagenome-assembled contigs in order to obtain the read count features described in Section 2.1. We use samtools v1.9 (Li *et al.*, 2009) to extract coverage and single nucleotide polymorphism (SNP) information from each bam file generated from the Bowtie2 mapping.

### 2.3 Model and optimization

DeepMAsED is a function *M*: ℝ^*L×p*^ → {0,1} providing a score *M*(*x*) representing the confidence that a contig x is misassembled. We denote by *y* ∈ {0,1} the label of the corresponding contig, where *y* = 1 if x is *extensively misassembled* according to MetaQUAST. The detailed architecture for DeepMAsED can be found in Table S2.

First, the input contig and associated count features are passed through a convolutional feature extractor *E*, whose output is a feature vector **h** = *E*(x). The hidden vector **h** is then passed through fully connected layers, resulting in a scalar *F*(**h**) ∈ ℝ. Finally, we compute the output scores *M*(x) = *σ*(*F*(**h**)) ∈ [0,1], where *σ* is the sigmoid function 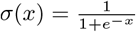.

The model is trained to minimize the weighted binary cross-entropy loss

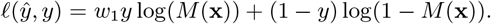

The contribution of the loss of positive instances is up-weighted by *w*_1_ = (*n* — *P*)/*P*, where *P* is the number of contigs labeled as misassembled in the training set. The loss function is optimized using Adam (Kingma and Ba, 2015) with mini-batches of size 64. We use batch-normalization (Ioffe and Szegedy, 2015) between convolutional layers and dropout (Srivastava *et al.*, 2014) with rate 0.5 between fully-connected layers.

**Figure 2:**
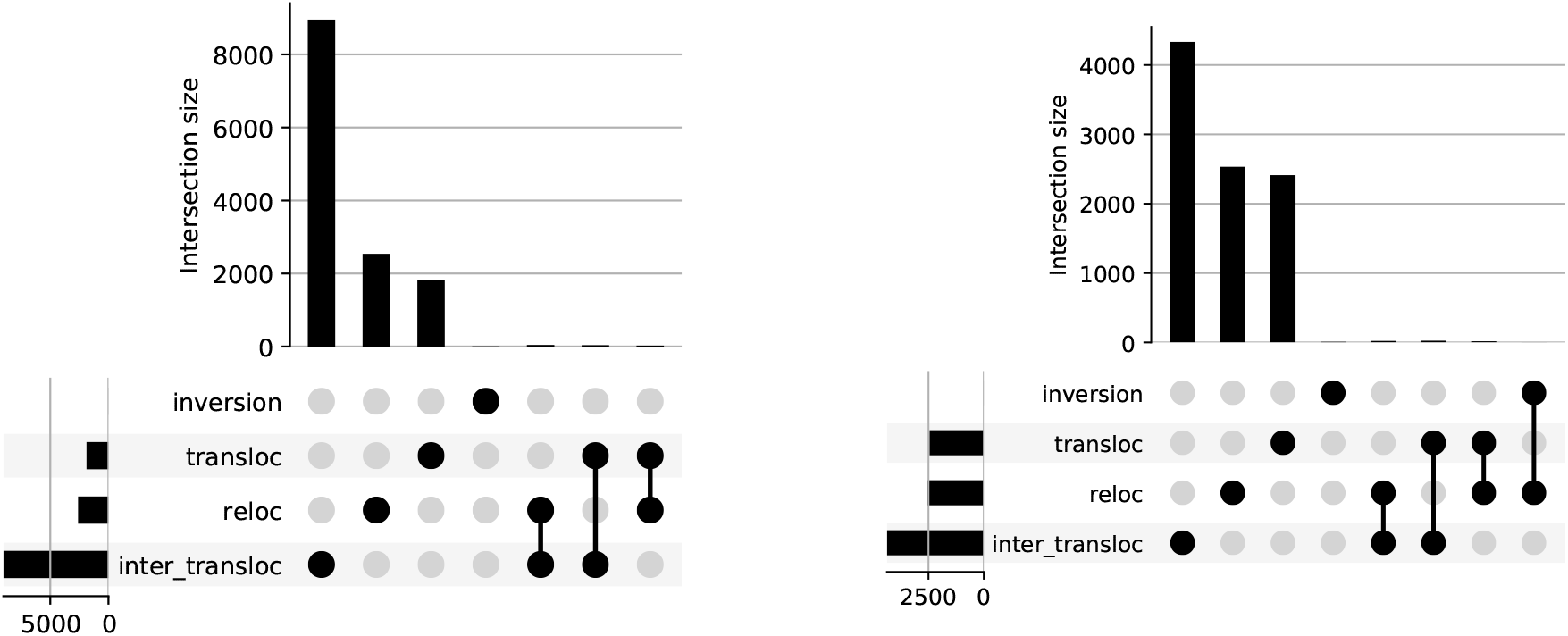
Distribution of errors found by metaQUAST (Mikheenko *et al.*, 2015) in the training set (left) and test set (right). The labels “inversion”, “transloc”, “reloc”, and “inter_transloc” stand for *inversion, translocation, relocation*, and *inter-genome translocation* respectively, as defined by metaQUAST.

Given the strong class imbalance, we select *average precision* as a validation metric, which corresponds to the area under the curve for the precision-recall curve. To encourage generalization to diverse data distributions, we train the model using 5-fold cross-validation over metagenomes as follows. Given 30 metagenomes available for training, we successively reserve 6 of them for computing the average precision, and the remaining 24 for training. This ensures that the validation data is drawn from metagenomes with different original abundances relative to the training data. Each of the five resulting models is trained over for 5 epochs over the data. The model is jointly trained on contigs assembled from MEGAHIT and MetaSPAdes.

The parameters leading to the highest average cross-validation precision are selected and are provided in the supplementary material. Finally, a model with the chosen parameters is trained on all the available training data from both assemblers for 10 epochs over the data.

### 2.4 Interpretability

Given their black-box nature, the interpretability of neural networks remains an open area of research (Gilpin *et al.*, 2018). A number of attempts has been made to provide feature importance scores for inputs to the network, that is, quantifying the impact that individual input features have on the prediction. We provide feature importance for DeepMAsED using DeepLIFT (Shrikumar *et al.*, 2017). Given an input contig, we use 10 references by using the di-nucleotide shuffling function from DeepLIFT.

### 2.5 Comparison to competing methods

Few methods are available for evaluating the quality of MAGs directly at the contig level without one or more reference genomes. Two predominant methods in this field, ALE (Clark *et al.*, 2013) and SuRankCo (Kuhring *et al.*, 2015), are no longer actively maintained. In particular, SuRankCo cannot be used at the time of this work due to software incompatibilities, so we could not compare against it.

ALE attributes a likelihood score to each individual position in the assembled sequence, not to a whole contig, and consists of four sub-scores: *depth, place, insert*, and *kMer* log-likelihoods. The tool utilizes strong probabilistic assumptions about the contig data such as independence of the errors quantified by each sub-score and also species-specific tetra-nucleotide frequencies.

In order to aggregate ALE scores into a contig score, we proceed similarly to Kuhring *et al.* (2015). We select a threshold for each sub-score and count the number of positions in the contig whose likelihood value is smaller than the threshold. The total count is normalized by length and the number of sub-scores to obtain a score *s* ∈ [0,1] for a given contig.

The value of the four thresholds is obtained via parameter search on the training data: for each assembler and each out the 30 generated metagenomes, we compute the average precision given the input thresholds. We then compute the mean average precision for all 30 metagenomes, resulting in an average precision scores of *AP_MH_* and *AP_MS_* for MEGAHIT and MetaSPAdes, respectively for each combination of thresholds. Since DeepMAsED is trained on contigs from both assemblers, we then compute for each threshold combination the average 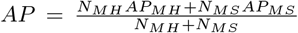, where *N_MH_* and *N_MS_* are the total number of contigs from each assembler. We select the thresholds leading to the highest average precision in this fashion. The thresholds considered can be found in Table S3.

Since CheckM (Parks *et al.*, 2015) is the most widely used measure of MAG assembly quality, we compare our predictions with both CheckM-estimated completeness and contamination on data from Almeida *et al.* (2019) and Pasolli *et al.* (2019).

## 3 Results

### DeepMAsED generalizes to new metagenomes and outperforms ALE

DeepMAsED generalizes to the test data, see Figure 3. This happens despite the mismatch in misassembly class distribution between the training and test data (Figure 2). Moreover, the genomes from the training and test datasets do not overlap, suggesting that DeepMAsED generalizes well to novel genomes.

**Figure 3:**
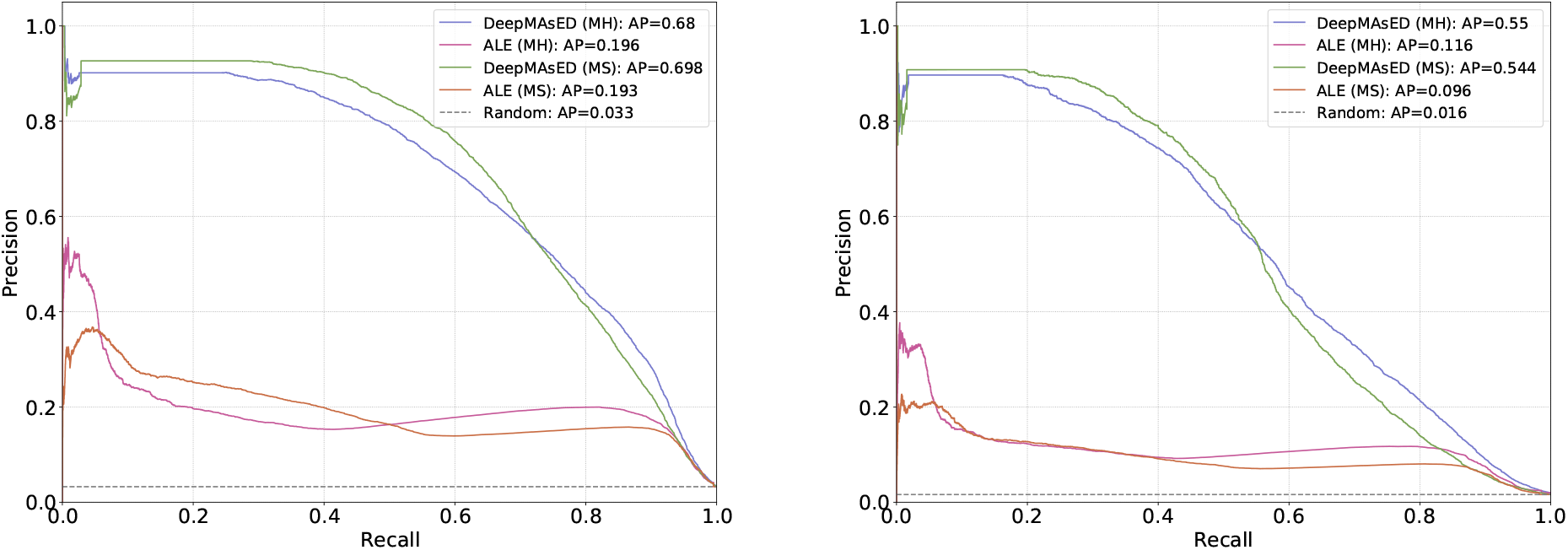
Average precision on the synthetic datasets on the training set (left) and test set (right). DeepMAsED generalizes to the test data and significantly outperforms ALE and random prediction. On the test data, a threshold achieving 25% recall reaches 87% precision and a threshold achieving 50% recall reaches 62% precision. We use both these thresholds for computing *conservative* and *flexible* estimates on the real world datasets.

On the test data, DeepMAsED achieves a substantially higher precision relative to ALE for both assemblers at any recall threshold below 90%. For instance, precision is close to 90% at 25% recall for DeepMAsED, while ALE only achieves close to 10% precision at the same recall. Both methods significantly outperform random prediction, which achieves low precision due to class imbalance. While achieving high precision at high recall remains an open problem, DeepMAsED provides a conservative method to identify misassembled contigs.

### DeepMAsED predicts a substantial number of misassemblies in benchmark datasets

We applied DeepMAsED on two recent large-scale metagenome assembly studies, Almeida *et al.* (2019) and Pasolli *et al.* (2019), in which 92 143 and 154723 MAGs were generated from the assembly of thousands of human metagenome samples. Both studies utilized CheckM estimations of MAG completeness and contamination. Given the very large numbers of metagenomes used in Almeida *et al.* (2019) and Pasolli *et al.* (2019), we sub-sampled the datasets to 1438 and 509 MAGs assembled from 133 and 51 metagenomes, respectively.

We select two thresholds for DeepMAsED prediction: a *conservative* threshold achieving 25% recall on the test data, and a *flexible* estimate achieving 50% recall on the test data. These thresholds achieve precisions of 87% and 62% respectively, see Figure 3 (right).

The flexible estimate finds 3564 misassembled contigs in Almeida *et al.* (2019) and 1136 in Pasolli *et al.* (2019), see Table 1. This represents close to 5% misassembled contigs in both datasets. ALE identifies significantly more misassembled contigs, but the false positive rate is likely high, given the low precision achieved on the test data.

**Table 1:** Number of contigs classified as misassembled. The conservative and flexible estimates correspond to recalls of 25% and 50% respectively on the test set. At these recalls, DeepMAsED achieves precisions of 87% and 62% respectively, while ALE achieves 21% and 15%. Only contigs and MAGs present in the output of ALE were used, resulting in 21034 contigs for Pasolli *et al.* (2019) and 71825 for Almeida *et al.* (2019). Values in parentheses are percentages of all contigs used. DeepMAsED predicts 5% of misassembled contigs at 50% recall.

| Method | Conservative | Flexible |
| --- | --- | --- |
| DeepMAsED (Almeida) | 30 (0.05%) | 3 564 (5.0%) |
| ALE (Almeida) | 3 617 (5.0%) | 37 020 (51.5%) |
| DeepMAsED (Pasolli) | 12 (0.05%) | 1 136 (5.4%) |
| ALE (Pasolli) | 1 704 (8.1%) | 12 685 (60.3%) |

### CheckM scores are poor proxies for DeepMAsED predictions

Given that CheckM has become the *de facto* standard for determining MAG quality (usually based on a cutoff of ≥ 50% completeness and < 5% contamination), we sought to determine how well CheckM quality estimates correlate with the prevalence of contig misassemblies, as identified by DeepMAsED. Both contamination and completeness scores from CheckM are poor indicators of DeepMAsED scores, see Figure 4 for contigs from Almeida *et al.* (2019). Scores for contigs from Pasolli *et al.* (2019) are provided in Figure S4.

**Figure 4:**
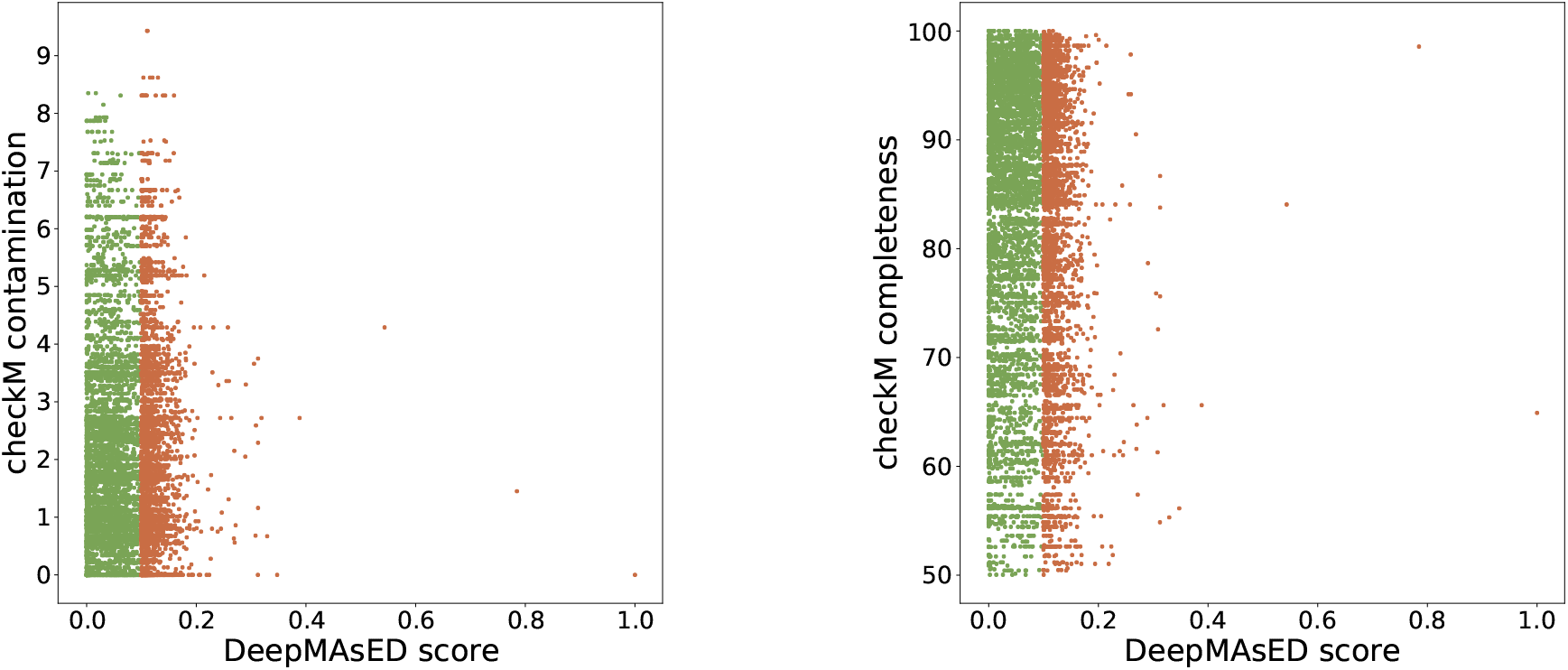
CheckM contamination (left) and completeness (right) for each contig in Almeida et *al.* (2019) against the corresponding DeepMAsED score. Points in brown are classified as misassembled with a flexible estimate at 50% recall. Both CheckM completeness and contamination are poor proxies for DeepMAsED quality predictions. Given the large number of contigs, contigs in green were subsampled to 5 000 points to reduce overplotting.

### DeepLIFT highlights critical features for high misassembly scores

DeepLIFT scores on individual contigs spike mainly in two scenarios. First, input locations with low coverage and an increased number of SNPs, see Figure 5 (top). Second, input locations with high coverage but a high number of SNPs, see Figure 5 (bottom). Further examples can be found in Figures S5 and S6. These results suggest that DeepLIFT, when used with DeepMAsED, can help to interpret why contigs are classified as misassemblies and to identify specific contig regions where misassemblies have likely occurred.

**Figure 5:**
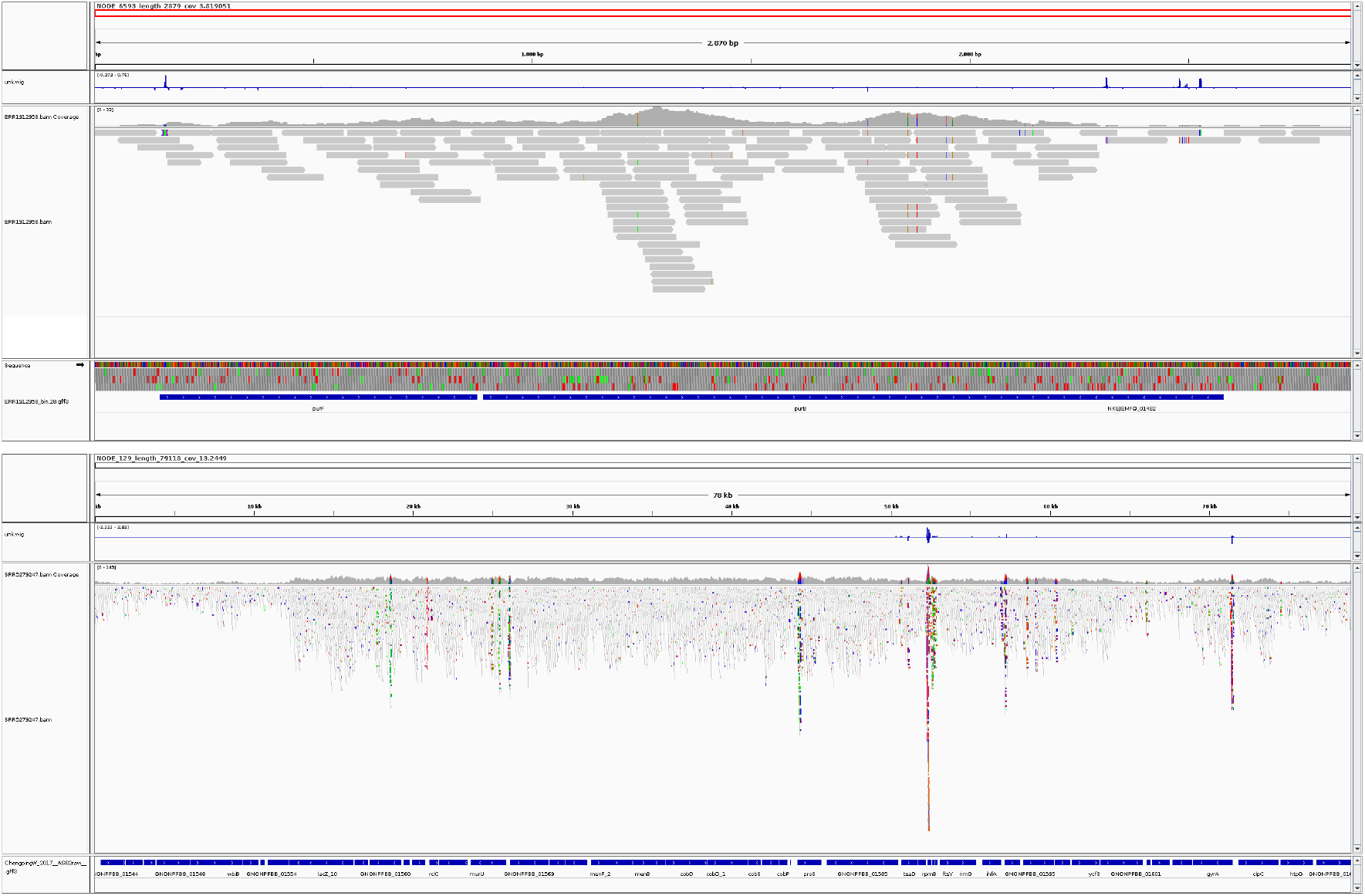
Visualization of the DeepLIFT (feature importance) output on two contigs from Almeida *et al.* (2019) and Pasolli *et al.* (2019), `NODE_6593_length_2879_cov_3.819051` and `NODE_129_length_79118_cov_13.2449`, which aims to display the effect on different input locations on the predicted score. The peaks of DeepLIFT activations can be seen on the blue line on top of the coverage information. The bottom figure shows high activation in a zone with large coverage and a large number of SNPs. On the contrary, the top figure display high activations on zones of low coverage. For each plot, the sections are: contig location, deepLIFT values, coverage, reads mapped with SNPs shown in various colors, and genes.

### Performance

We evaluate DeepMAsED using an NVIDIA K80 GPU. We benchmark performance by computing average time per mini-batch over 100 mini-batches of size 64 over the training data. The average time per mini-batch is 0.084 ± 0.00 seconds, or 42 418 contigs of size 10 000 per minute. Using only an Intel Xeon CPU E5-2680 v4 at 2.40GHz CPU, evaluation is slower, at 0.75 ± 0.05 seconds per mini-batch, or 4 587 contigs per minute. The final model has 102 627 trainable parameters, requiring a modest 1.3 MB of storage.

## 4 Discussion

### DeepMAsED generalizes to a large number of research projects

The results in Section 3 suggest that DeepMAsED generalizes to a large variety of genomes and metagenome compositions for two widely used assemblers. In order for researchers to deploy DeepMAsED on their assemblies, input features must be computed. This requires mapping reads to contigs, but this is a usual step in other downstream applications such as estimating MAG relative abundances. Moreover, we provide code to compute the input features from raw BAM files and weight matrices from the fully trained system, which then allows researchers to easily assess the quality of their assembled contigs. Misassembly classification is rapid and should scale to millions of contigs when using a GPU.

We also provide code to generate large amounts of realistic data in a straight forward manner. This allows researchers to train and evaluate new models from scratch with larger or more specific reference genome datasets (e.g., using biome-specific microbes or virus genomes).

### Future steps toward improving performance

A first potential direction for improving DeepMAsED performance is increasing the amount of training data. This could be achieved in several ways such as increasing the number of genomes, the number of simulated metagenomes, and/or the corresponding sequencing depth. More metagenome assemblers could be included to increase generalization, but few show the scalability, accuracy, and popularity of MEGAHIT and MetaSPAdes (Wang *et al.*, 2019).

In another direction, DeepMAsED could be trained on long read or hybrid short-long read metagenome assemblies. Especially for assemblies solely utilizing error-prone long reads (e.g., Oxford Nanopore reads), the misassembly error profile may substantially differ from the short read assemblies evaluated in this work (Nicholls *et al.*, 2019).

Performance could be potentially improved by changing how long contigs are evaluated, which is currently batched in non-overlapping windows of fixed sized. A sliding window may improve performance, but possibly at a high computational cost. Also, we note that our currently trained model that we provide should only be used with contigs longer than 1 000 bp, as shorter contigs were filtered during training.

We could also enrich the features present at each location in the input contigs. One possibility is adding gene annotations, with the assumption that misassemblies could interfere with gene calling (i.e., no annotation) or produce poor annotations (e.g., genes of unknown function). This option adds significant overhead, since gene calling and annotation are computationally expensive. Another possibility is adding taxonomy information at each position by majority voting from a sliding window taxonomy, or using the entropy of the taxonomy sliding window. High entropy could signal a rapidly changing taxonomy in the region, which may be a result of misassembly-generated chimerism. The main drawback of taxonomy features, in addition to computational costs, is the quality of the reference taxonomies themselves. This is particularly relevant in the context of novel genome assemblies, which can lack close relatives in reference taxonomies. One could also add ALE features as inputs to DeepMAsED, but this would add strong assumptions that may not hold for novel genomes.

Finally, we could improve DeepMAsED by providing more fine-grained predictions for the type of misassembly present (e.g., inversion or relocation) or identifying the location of the misassembly directly. This would be of particular value for long contigs in which a large portion of the contig may still be correctly assembled.

## 5 Conclusion

DeepMAsED is a generally applicable tool for evaluating metagenome assembly accuracy. It demonstrates that deep learning is a tractable solution for reference-free identification of metagenome-assembled contig misassembly and provides a benchmark from which to improve upon. While there is room for improving the accuracy of DeepMAsED, we note that regardless of the misasssembly classification threshold used, the researcher can use DeepLIFT, ALE, metaQUAST or other tools to investigate the subset of contigs tentatively identified as misassemblies by DeepMAsED. This hybrid automated-manual curation approach should greatly reduce the workload relative to fine-grained checking of all contigs individually.

## 6 Acknowledgements

This work was supported by the Max Planck Society. We thank Guillermo Luque for helpful discussions on this work.

## Footnotes

1 *mased*: a Middle English term for “misled”

## Notes

https://github.com/leylabmpi/DeepMAsED

