## Supplemental Materials for "DeepMAsED: Evaluating the quality of metagenomic assemblies"

**Supplementary Material for: *DeepMAsED: Evaluating the quality of metagenomic assemblies***

### 1 Network architecture and parameters

Parameter search for DeepMAsED was done via cross-validation on the parameters in Table S1. The best validation accuracy was obtained by $n_f = 8$, $n_c = 5$, $n_h = 50$ and $p = 50$. Moreover, the initial learning rate for Adam was 0.0001. We selected a 0.5 dropout rate by observing that smaller values systematically led to over-fitting.

| **Parameter** | **Values** |
|---|---|
| Number of initial filters $n_f$ | $\{8, 16\}$ |
| Number of convolutional layers $n_c$ | $\{3, 4, 5\}$ |
| Number of hidden units in FC layers $n_h$ | $\{30, 50, 70, 256\}$ |
| Size of pooling window $p$ | $\{30, 50, 70\}$ |

Table S1: Parameters tested during DeepMAsED training.

The final network architecture for DeepMAsED is provided in Table S2.

| DeepMAsED Layers | Output size |
|---|---|
| Input | $10\,000 \times 11 \times 1$ |
| Conv $2 \times 11 \times 8$, stride 1 + ReLU + BN | $10\,000 \times 1 \times 8$ |
| Conv $2 \times 1 \times 16$, stride 2 + ReLU + BN | $5\,000 \times 1 \times 16$ |
| Conv $2 \times 1 \times 32$, stride 2 + ReLU + BN | $2\,500 \times 1 \times 32$ |
| Conv $2 \times 1 \times 64$, stride 2 + ReLU+ BN | $1\,250 \times 1 \times 64$ |
| Conv $2 \times 1 \times 64$, stride 2 + ReLU+ BN | $625 \times 1 \times 128$ |
| Average Pooling $50 \times 1$ | $12 \times 1 \times 128$ |
| Flatten | $1\,538 \times 1$ |
| FC 50 + ReLU + Dropout 0.5 | $50 \times 1$ |
| FC 50 + ReLU + Dropout 0.5 | $50 \times 1$ |
| FC 50 + sigmoid + Dropout 0.5 | $1 \times 1$ |

Table S2: Architecture for DeepMAsED. **BN** stands for Batch Norm (Ioffe and Szegedy, 2015), **FC** $\mathbf{n}_\ell$ for fully connected with $n_\ell$ units. **Conv** $\mathbf{v} \times \mathbf{w} \times \mathbf{f}$ stands for a 2d convolutional layer with kernel size $(v, w)$ and $n_f$ filters. **Dropout** (Srivastava *et al.*, 2014) **r** stands for dropout with rate $r$. The total number of trainable parameters is 102 627.

### 2 Parameter search for ALE

The parameters tested as thresholds for the four ALE scores are provided in Table S3. The parameters leading to the best average precision are

$$d = -21,\ p = -2,\ i = 2,\ k = 2.$$

| Parameter | Values |
|---|---|
| Depth $d$ | $\{-21, -19, \ldots, 0, 2\}$ |
| Place $p$ | $\{-9, -7, \ldots, 0, 2\}$ |
| Insert $i$ | $\{-9, -7, \ldots, 0, 2\}$ |
| k-Mer $k$ | $\{-9, -7, \ldots, 0, 2\}$ |

Table S3: Parameters tested as ALE thresholds.

### 3 Similarity of training and test genomes

The distribution of contig lengths for the generated training and test datasets can be found in Figure S1. Moreover, the ANI similarity between the genomes in the training and test datasets can be found in Figure S2.

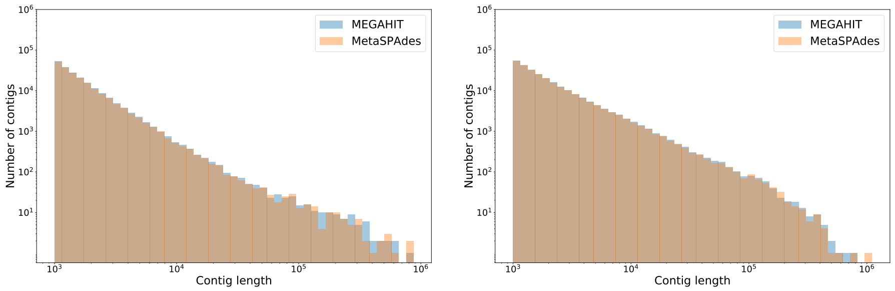


Figure S1: Length distribution of the assembled contigs in the training (left) and test sets (right) in logarithmic scale, obtained both from MEGAHIT and MetaSPAdes.

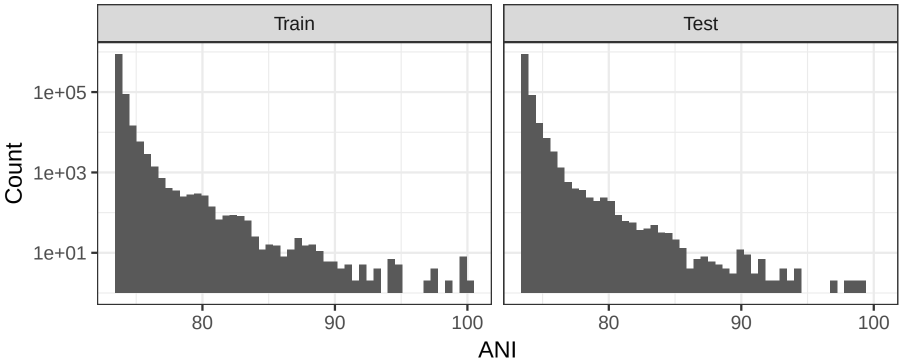


Figure S2: ANI between genomes in the training (left) and test (right) datasets.

### 4 CheckM scores for the train/test genomes

The CheckM completeness and contamination scores for the genomes in the training and test datasets is given in Figure S3.

### 5 CheckM scores for contigs from Pasolli *et al.* (2019)

CheckM completeness and contamination scores for contigs from Pasolli *et al.* (2019) can be found in Figure S4.

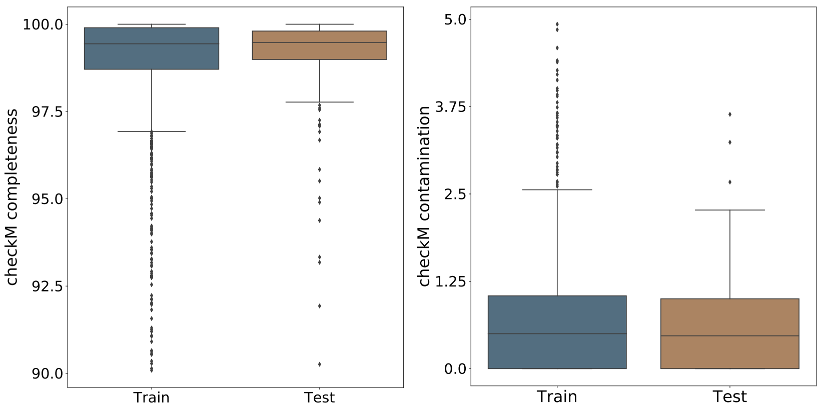


Figure S3: CheckM contamination (left) and completeness (right) in the training and test datasets.

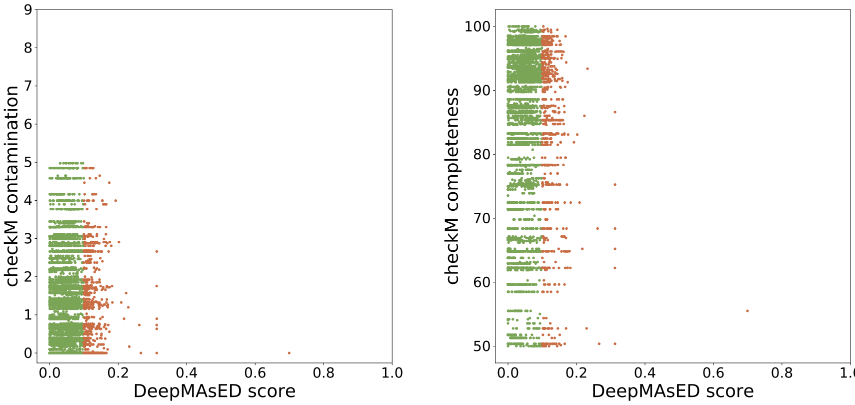


Figure S4: CheckM contamination (left) and Completeness (right) for each contig in Pasolli *et al.* (2019) against the corresponding DeepMAsED score. Points in brown are classified as misassembled with a flexible estimate at 50% recall. Both checkM completeness and contamination are poor proxies for DeepMAsED quality predictions. Given the large number of contigs, contigs in green were subsampled to 5 000 points to reduce overplotting.

### 6 Further DeepLIFT examples

We provide further examples of DeepLIFT feature importance visualizations in Figures S5 and S6.

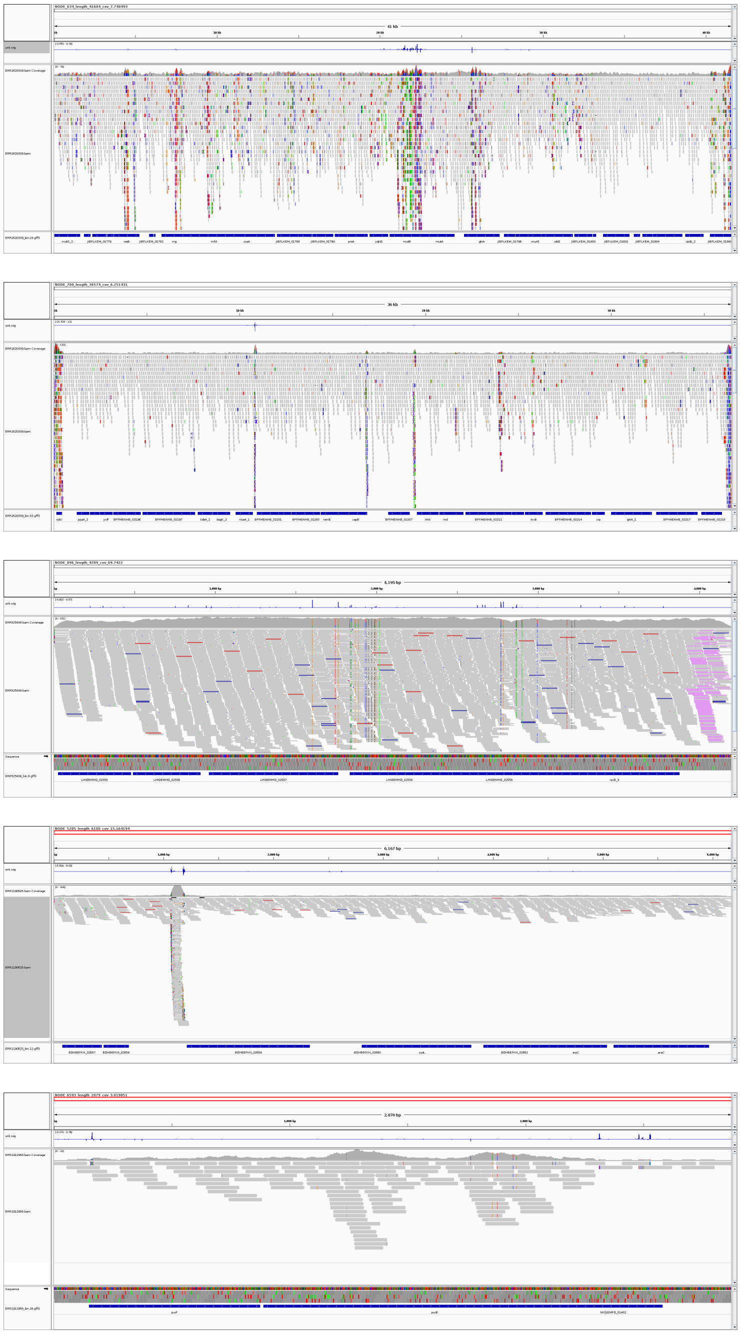

Figure S5: Visualization of the DeepLIFT (feature importance) output for contigs from the dataset in Almeida *et al.* (2019).

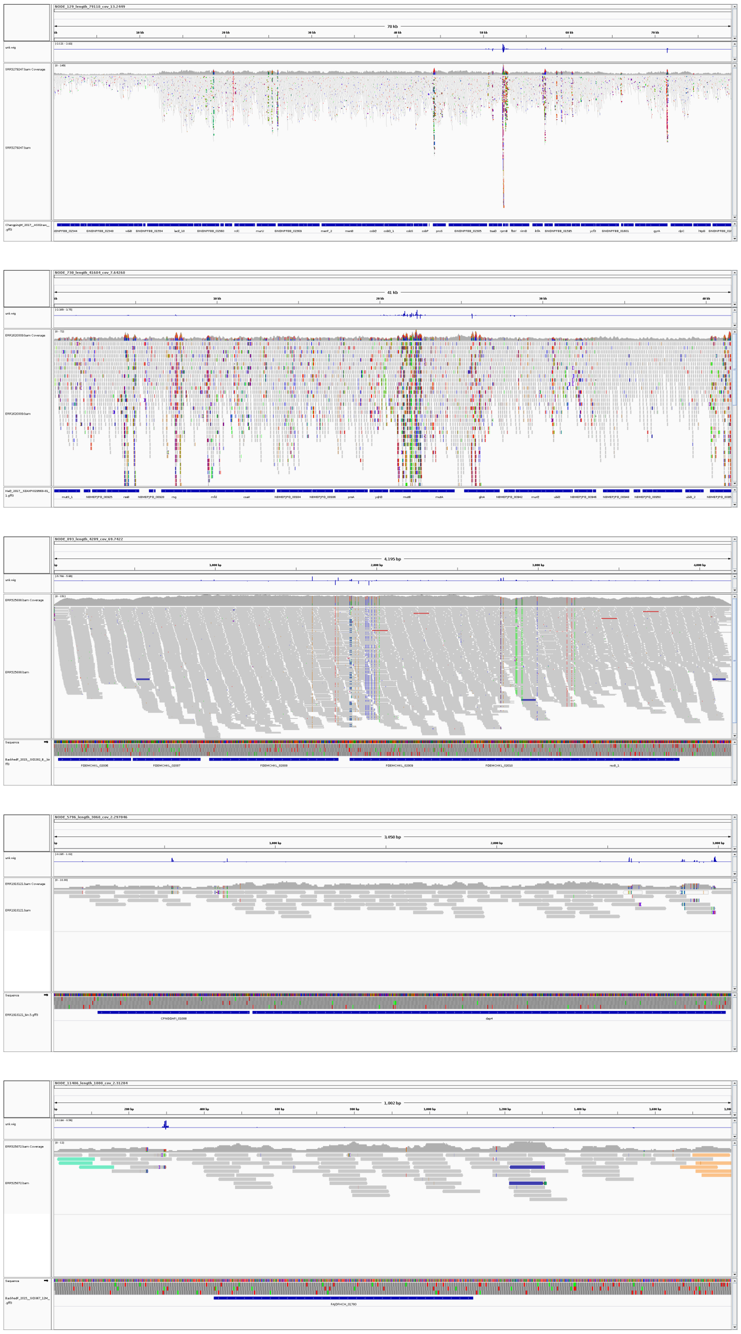

Figure S6: Visualization of the DeepLIFT (feature importance) output for contigs from the dataset in Pasolli *et al.* (2019).
